# Major QTLs for resistance to early and late leaf spot diseases are identified on chromosomes 3 and 5 in peanut (*Arachis hypogaea*)

**DOI:** 10.1101/567206

**Authors:** Ye Chu, Peng Chee, Albert Culbreath, Tom G. Isleib, Corley C. Holbrook, Peggy Ozias-Akins

## Abstract

Early and late leaf spots are the major foliar diseases of peanut responsible for severely decreased yield in the absence of intensive fungicide spray programs. Pyramiding host resistance to leaf spots in elite cultivars is a sustainable solution to mitigate the diseases. In order to determine the genetic control of leaf spot disease resistance in peanut, a recombinant inbred line population (Florida-07 x GP-NC WS16) segregating for resistance to both diseases was used to construct a SNP-based linkage map consisting of 855 loci. QTL mapping revealed three resistance QTLs for late leaf spot *qLLSA05* (phenotypic variation explained, PVE=7-10%), *qLLSB03* (PVE=5-7%), and *qLLSB05* (PVE=15-41%) that were consistently expressed over multi-year analysis. Two QTL, *qLLSA05* and *qLLSB05*, confirmed our previously published QTL-seq results. For early leaf spot, three resistance QTLs were identified in multiple years, two on chromosome A03 (PVE=8-12%) and one on chromosome B03 (PVE=13-20%), with the locus *qELSA03_1.1* coinciding with the previously published genomic region for LLS resistance in GPBD4. Comparative analysis of the genomic regions spanning the QTLs suggests that resistance to early and late leaf spots are largely genetically independent. In addition, QTL analysis on yield showed that the presence of resistance allele in *qLLSB03* and *qLLSB05* loci might result in protection from yield loss caused by LLS disease damage. Finally, post hoc analysis of the RIL subpopulation that was not utilized in the QTL mapping revealed that the flanking markers for these QTLs can successfully select for resistant and susceptible lines, confirming the effectiveness of pyramiding these resistance loci to improve host-plant resistance in peanut breeding programs using marker-assisted selection.

## Introduction

Peanut (*Arachis hypogaea* L.) is an important row crop rich in oil, protein, vitamins and other micronutrients (Settaluri et al., 2012). World peanut production exceeded 100 thousand metric tons in year 2016. The United States is the most efficient peanut producing country and accounts for 6% of world production (http://www.fao.org). Peanut production is threatened by multiple biotic stresses, of which the two foliar fungal diseases, early leaf spot (ELS) caused by *Passalora arachidicola* (*Hori*) U. Braun (syn. *Cercospora arachidicola*) and late leaf spot (LLS) caused by *Nothopassalora personata* (Berk.& M.A. Curtis) U. Braun, C. Nakash., Videira & Crous (syn. *Cercosporidium personatum*), predominate. Both fungal diseases produce lesions (up to 1 cm in diameter) on peanut leaves, petioles, stems and pegs (McDonald et al., 1985). Shedding of infected leaves upon disease progression can lead to complete defoliation in susceptible genotypes and up to 70% yield loss (Singh et al., 2011). In the U.S., the most common practice to control both diseases is by frequent fungicide applications; therefore, it is not surprising that fungicide sprays to control leaf spot diseases incur the highest cost in disease management (Woodward et al., 2014). Developing and planting ELS and LLS resistant peanut cultivars should reduce the cost of peanut production while simultaneously mitigating the environmental footprint through reduction in pollution from fungicides.

The physical appearance of early and late leaf spot lesions is similar but the two diseases can be accurately diagnosed for their causal agent by the position of sporulation in the field (McDonald et al., 1985) (Figure 1). Sporulation of *P. arachidicola* predominately occurs on the adaxial side of peanut leaves, thus lesions from ELS are often dark brown on the adaxial (upper) surface of a peanut leaflet and coupled with a chlorotic halo (Figure 1A). Lesions from LLS, on the other hand are black in color (Figure 1B) and N. *personata* produces spores mostly on the abaxial (under) side of the leaf. Additionally, as the disease name indicates, ELS occurs earlier in the growing season than LLS. In the southeastern U.S. peanut growing region, ELS occurs between June and August whereas LLS emerges around mid- to late-August. However, the epidemic of the two fungal diseases varies across peanut growing regions. In recent years, LLS has increased in prevalence in the southeastern region (Cantonwine et al., 2008) and has been reported as the dominant foliar disease in Georgia, the number one peanut producing state in the U.S. (Fulmer, 2017). In the Virginia-Carolina region, however, ELS is the predominant leaf spot disease which led to the field selection of ELS-resistant lines, including GP-NC WS16, through introgression from wild species (Stalker, 2017).

**Figure 1.**
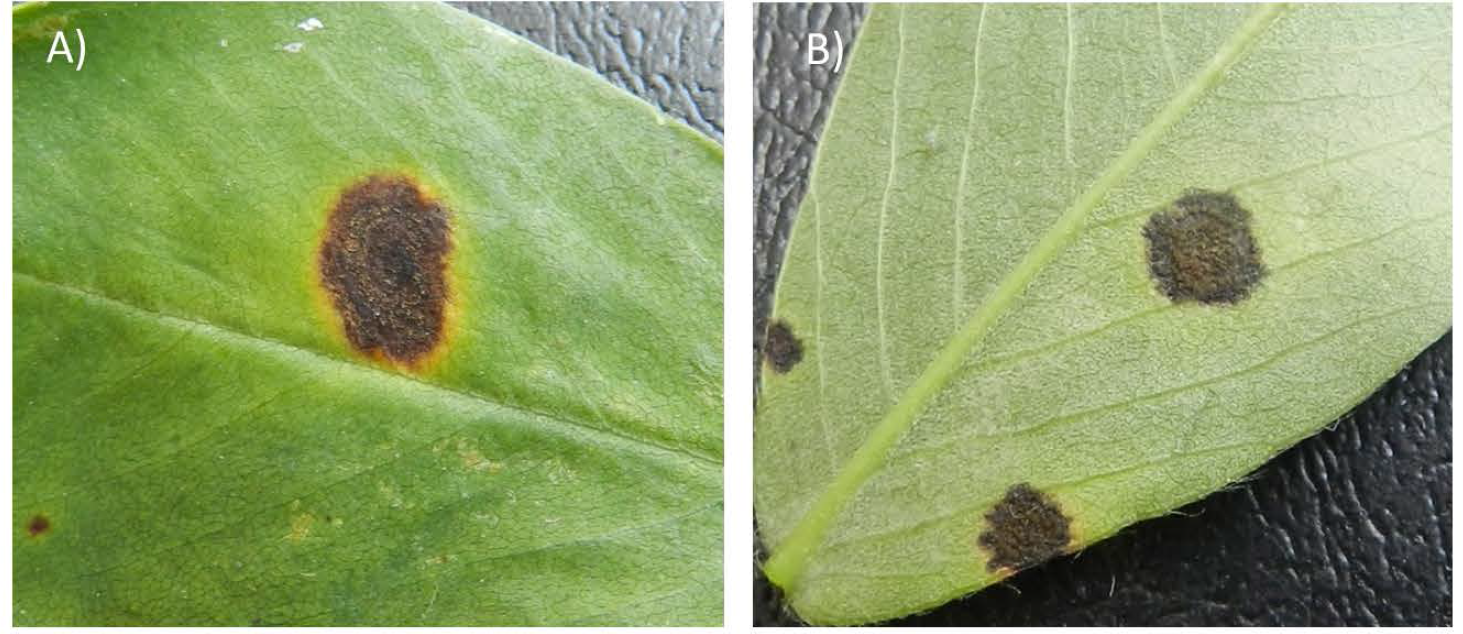
Early leaf spot lesion (A) and late leaf spot lesion (B) formed on peanut leaves. The early leaf spot lesion typically causes a yellow halo surrounding the lesion whereas the late leaf spot lesion often lacks the halo (B). However, the most important criterion to distinguish the diseases lies in the location of sporulation. Early leaf spot sporulates predominantly on the adaxial (upper) side of a leaf, whereas late leaf spot sporulates on the abaxial (lower) side of a leaf.

Cultivated peanut is an allotetraploid (AABB genome) with a genome size of 2.7 Gb (Dhillon et al., 1980; Bertioli et al., 2016; Nascimento et al., 2018). It’s inbred nature, recent domestication, perhaps single polyploidization event, and the presence of crossing barriers between the cultivated species lines and its wild diploid relatives contribute to the paucity of genetic polymorphism within this species. Earlier marker types such as restriction fragment length polymorphisms (RFLP), amplified fragment length polymorphisms (AFLP), simple sequence repeats (SSR), and cleaved amplified polymorphic sequences (CAPS) had limited ability to detect genetic polymorphism within the cultivated peanut germplasm (Pandey et al., 2012). Recent advancements in next generation sequencing and bioinformatics pipelines for SNP calling (Clevenger et al., 2015; Clevenger and Ozias-Akins, 2015) have made SNP markers accessible for peanut genetic mapping. Double-digested RAD-seq was applied on a mapping population and produced the first SNP-based peanut genetic map with 1,685 loci covering a total map distance of 1,446 cM (Zhou et al., 2014). More recently, a large SNP array with 58K loci was developed providing a high throughput genotyping platform for the peanut genetics research community (Clevenger et al., 2017b).

Several sources of host resistance to leaf spot diseases have been reported previously. Among 500 U.S. peanut plant introductions (PI) tested, only 33 were reported to have partial resistance to LLS (Anderson et al., 1993). The LLS resistant line PI 203396 was used to develop the leaf spot resistant cultivars Georganic (Holbrook and Culbreath, 2008) and Tifrunner (Holbrook and Culbreath, 2007). Serendipitously, this PI also introduced resistance to tomato spotted wilt virus (TSWV) (Gorbet et al., 1987; Clevenger et al., 2017b). NC 94022 is a TSWV resistant breeding line derived from PI 576638, a *hirsuta* type of peanut (Culbreath et al., 2005), which was used as the LLS resistant parent for a mapping population (Khera et al., 2016). In addition, a Texas breeding line Tx964117 was reported to have resistance to both ELS and LLS (Liang et al., 2017). Besides these resistance sources from cultivated peanut, wild peanut relative *A. cardenasii* was found to harbor strong resistance to both ELS and LLS (Abdou et al., 1974). Introgression of *A. cardenasii* chromatin accomplished through a synthetic hexaploid bridge has led to the release of multiple ELS resistant lines including the male parent of our mapping population GP-NC WS 16 (Stalker and Beute, 1993; Stalker et al., 2002; Tallury et al., 2014). The same resistance source was sent to ICRISAT, India in the 1980s and used in breeding programs resulting in the release of GPBD 4 and ICGV86699 (Reddy et al., 1996; Stalker, 2017).

Several leaf spot resistant sources have been utilized in genetic mapping to identify QTLs for resistance to ELS and LLS. For example, genetic mapping of LLS resistance using the GPBD 4 derived recombinant inbred line (RIL) population yielded 11 QTLs on a genetic map with 56 SSR marker loci (Khedikar et al., 2010) and 28 QTLs on an improved SSR map with 260 loci (Sujay et al., 2012). Further mapping with 139 additional SSR and transposable element markers revealed two major QTL regions for LLS resistance on chromosomes A03 and B10 (Kolekar et al., 2016). Further QTL-seq analysis by whole genome resequencing identified a salient wild introgression located at 131.6 to 134.7 Mb of chromosome A03 (Pandey et al., 2017a). One SNP marker within this region was converted to a gel-based dominant marker for marker-assisted breeding. Genetic mapping with the Zhonghua 5 by ICGV 86699 population, which consists of 1,685 SNP loci from double-digested RAD seq (Zhou et al., 2016), detected 20 LLS QTLs with five out of six major QTLs located on chromosome B06. Unfortunately the genome positions of these QTLs were not available for comparison since the physical position was not reported. In the Tifrunner x GT-C20 population, 37 LLS-resistance QTLs were identified using a genetic map constructed from a F2 population with 318 marker loci (Wang et al., 2013). A slightly denser map was later developed using an F_8:9_ RIL population of the same cross with 418 loci on which 9 and 22 QTLs for ELS and LLS, respectively, were identified (Pandey et al., 2017b). In the SunOleic-97R x NC 94022 population, 22 QTLs for ELS and 20 for LLS were reported using a SSR-based map consisting of 248 loci (Khera et al., 2016). Finally, in the Tamrun OL07 x Tx964117 population, a 1211 loci SNP map was developed from double-digested RAD-seq markers and six QTLs contributed by the ELS resistant parent Tx964117 were identified on chromosomes A02, B04, B06, B09 and B10 (Liang et al., 2017). These studies suggest that different genetic sources of leaf spot resistance may harbor different genes/alleles that will be useful in breeding.

In this study, a RIL population developed from the cross of Florida-07 x GP-NC WS 16 was used to map QTLs associated with ELS and LLS resistance. This population was developed as part of a nested association mapping population (Holbrook et al., 2013). The female parent Florida-07 (Gorbet and Tillman, 2008) is a high oleic cultivar susceptible to LLS while the male parent, GP-NC WS 16, is a germplasm line derived from the same group of interspecific breeding materials with introgression from *A. cardenasii* as the ICRISAT lines GPBD 4 and ICGV86699 (Tallury et al., 2014; Stalker, 2017). GP-NC WS 16 was released by North Carolina State University for possessing multiple disease resistances including ELS. In a previous study, QTL-seq analysis was performed on LLS resistant and susceptible bulks from this population and identified three genomic regions on A05, B03, and B05 contributing to LLS resistance (Clevenger et al., 2018). Building on our previous work, a SNP-based linkage map using the Axiom_*Arachis* 58K SNP array was constructed for this population. QTL analyses confirmed the LLS QTLs identified by QTL-seq, more precisely delineated the positions of the three QTLs, and showed that these regions also are associated with yield. Furthermore, QTL mapping identified major QTLs conditioning ELS resistance. Interaction of QTL regions for ELS and LLS are discussed.

## Materials and methods

### Recombinant inbred line population (RIL) and phenotyping

The RIL population utilized in this study was derived from a cross of Florida-07 and GP-NC WS 16 (greenhouse, 2012). Hybridity of the F_1_ plants was confirmed by genotyping with SSR marker GM1555 (Guo et al., 2012). Half of the F_2_ population was field advanced in Georgia via single seed descent to the F_6_ generation where single plants from each line were randomly selected and seeds from each were bulked to yield 192 RILs (hereafter referred to as the GA subpopulation). The other half of the F_2_ population was advanced in the same manner in North Carolina yielding 191 RILs (NC subpopulation). The GA subpopulation was advanced one year ahead of the NC subpopulation and was extensively phenotyped for both early and late leaf spot diseases. The NC subpopulation was only phenotyped for early leaf spot in 2015.

The GA subpopulation together with the two parents was planted for LLS evaluation following a randomized complete block design with three replications at the University of Georgia, Tifton GA described previously (Clevenger et al., 2018). In order to more precisely measure the leaf spot disease progression after the emergence of disease symptoms, disease ratings were taken four times, at 10 to 14 days intervals. The Florida rating scale, which takes into account both lesion coverage and percent defoliation, was used (Knauft et al., 1988). The rating scale ranged from 1, indicating no disease, to 10, indicating total plant death. The area under the disease progress curve (AUDPC) was calculated for LLS disease ratings (Shaner and Finney, 1977). Plot yield was collected after these LLS field trials from 2012 to 2014 and expressed as gram/plot.

Field trials for ELS were conducted in 2013 and 2014 for the GA subpopulation at the NCDA Peanut Belt Research Station located at Lewiston-Woodville, NC State University, NC. In 2015, the NC subpopulation was phenotyped for ELS resistance. A 14×14 lattice design was used with three field replications. Each plot consisted of two rows of 3.7 m in length with 14 seeds per plot row planted at 12.7 cm spacing. Four disease ratings were taken with an interval of 8 to 10 days since the emergence of the disease for year 2013 and 2014 trials. Two disease ratings was taken at 140 and 147 day after planting in year 2015 trial. The disease ratings were taken using the Florida Scale as previously described. Area under the disease progress curve (AUDPC) was calculated for each RIL.

### Genotyping

Both the GA and NC subpopulations were genotyped using the Axiom_*Arachis* 58K SNP version 1 array, but only the GA subpopulation was genotyped with SSR markers and utilized for linkage map construction and QTL analyses. A total of 409 SSR markers evenly distributed across the peanut genome were screened for parental polymorphism, and 65 polymorphic SSR markers were used in genotyping. Six previously described KASPar markers (Clevenger et al., 2018) were also used to genotype the GA subpopulation. Finally, since Florida-07 is a high oleic variety, the ahFAD2B_hybprobe targeting the A^442^ insertional mutation on the *ahFAD2B* gene (Chu et al., 2011) was mapped on the GA subpopulation. DNA was extracted from leaf tissue collected from 10 to 15 plants per line using Qiagen Plant DNeasy kit (Qiagen, Germantown, MD). DNA quantification was performed with Quant-iT Picogreen dsDNA assay kit (Thermo Fisher Scientific, Waltham, MA). For SSR markers, all forward primers were labeled with one of the fluorescent dyes FAM, TAMRA or VIC. PCR amplifications for SSRs used a touchdown program, starting with 95 C for 5 min, followed by 6 cycles of 95 C for 30s, 64 C (minus 1 C/cycle) for 30s and 72 C for 30s; followed by 30 cycles of 95 C for 30s, 58 C for 30s and 72 C for 30s; final extension was performed at 72°C for 7 min. PCR products were submitted to the Georgia Genomics and Bioinformatics Core (University of Georgia, Athens) for fragment analysis. KASPar marker development was described previously (Clevenger et al., 2018). Thermal cycling was performed on the Roche LC480 (Roche, Basel, Switzerland) as follows: 95°C for 15 min, followed by 9 cycles of 94°C for 20s and 61°C for 60s, with the annealing temperature dropped at the rate of 0.6°C/cycle, followed by 27 cycles at 94°C for 10s and 55°C for 60s and 2 cycles of 94°C for 20 s and 57°C for 60s. Pre- or post-melt cycles were performed at 30°C for 1s and cooling to 25°C during plate reading. For SNP array genotyping, DNAs were diluted to 30 ng/ul and submitted to Affymetrix for genotyping on the Axiom_*Arachis* SNP array with 58K probes (Clevenger et al., 2017b). The SNP data were analyzed using the Axiom Analysis Suite (Thermo Fisher Scientific). All polymorphic SNP loci between the parental lines were visually inspected for signal clustering. Markers demonstrating ambiguity in clustering were excluded from further analysis. SSR marker names were given as marker ID starting with the prefix GM and a numeric number. The allele size of Florida-07 was given after the SSR marker ID separated by an underscore. For SNP markers, those from the SNP array were given the prefix AX-followed by a nine digit code assigned by the company, while those from KASPar assays were assigned to a subgenome (A or B) followed by marker number and physical position. The high resolution melt marker distinguishing high oleic from normal oleic acid trait is designated as ahFAD2B_hybprobe (Chu et al., 2011).

### Genetic map construction

Only the GA subpopulation consisting of 165 RILs was utilized for linkage map construction. In addition to the SNP markers, a small number of SSRs, KASPar and the functional marker *ahFAD2B* gene were included as anchor markers to provide alignments of common loci between the previously published maps (Moretzsohn et al., 2005; Moretzsohn et al., 2009; Guo et al., 2012) and the current SNP map. The goodness of fit to the expected 1:1 segregation ratio for each locus (*P* < 0.05) was tested by Pearson’s Chi-square test. Sixty-two loci severely distorted from the expected segregation ratio (P<0.0001) were excluded. Linkage map construction was performed with the Joinmap software version 4.1 (https://www.kyazma.nl) with the minimum logarithm of odds (LOD) of 6.0. Kosambi map function (Kosambi, 1944) was used to transform the recombinant ratio into genetic distance. The genome positions of SNP markers were used to designate the A and B subgenomes for linkage groups (LGs). For the GA subpopulation, 27 RILs with greater than 5% heterozygosity were excluded from the dataset, leaving 165 RILs for linkage map construction and QTL analysis. A genetic map was drawn with Mapchart software version 2.32 (Voorrips, 2002).

### Data analysis

Statistical analysis of phenotyping data was performed with SAS software version 9.4 (*SAS* Institute Inc., 2016). Univariate variance analysis was performed by GLM method and the variance components were determined by restricted maximum likelihood method (REML). The broad sense heritability was estimated according to the formula: H^2^=σ_g_^2^/(σ_g_^2^ + σ^2^_gxe_/n+ σ^2^_e_/nr), where σ_g_^2^ was the genetic variance component among the RILs, σ^2^_gxe_ was the RIL x environment interaction variance component and σ^2^_e_ was the residual component, *n* was the number of environments and *r* was the number of replications (Hallauer and Miranda, 1988). Normality of data distribution was tested by the Shapiro test. Pearson correlation analysis was performed using the Proc corr procedure of SAS software.

QTL analysis was performed with the QTL cartographer 2.5 (Wang et al., 2005). Composite Interval Mapping (CIM) was performed to detect marker-trait association, using 1000 permutations to estimate the Likelihood Ratio (LR) threshold values (α = 0.05) to declare the presence of QTLs. The CIM analysis was performed at 1 cM walk speed in a 5 cM window by forward stepwise regression. QTL analysis was performed on the ELS, LLS and yield datasets from individual years separately. Since there were high positive correlations among the years and low genotype x year interactions for both disease traits, multiple years of data for each trait were combined by calculating their means and used for QTL analysis.

For those QTLs identified by only one year of disease rating, disease ratings collected at each time point of that year were analyzed with the same program. QTLs were reported when the AUDPC-derived QTLs were supported by single time point analysis. QTLs are designated following conventional nomenclature with the initial letter *q* followed by the trait name, linkage group and a numeric number indicating the number of QTL identified on the same LG.

In order to determine if there is significant wild introgression from *A. cardenasii* preserved in GP-NC WS 16, paired-end reads of GP-NC WS 16 (Clevenger et al., 2017b) and *A. cardenasii* were aligned with the *A. duranensis* reference genome (Bertioli et al., 2016). Diagnostic SNPs for alien introgression were identified by the Intromap pipeline (Clevenger et al., 2017a).

#### Validation of ELS QTLs in NC subpopulation

Flanking markers for each of the ELS QTLs were used to separate the NC subpopulation into two groups based on those possessing the Florida-07 or the GP-NC WS 16 alleles. Year 2015 ELS phenotyping data from the two groups were compared statistically using Student’s t-test. Statistical significance was declared at p-value <0.05.

#### Identification of candidate genes within the QTL intervals

In order to identify candidate genes potentially regulating disease resistance against the fungal pathogens, gene models were retrieved between the physical intervals of the consistent QTLs given in Table 2 from the *Arachis duranensis* and *Arachis ipaensis* genomes (peanutbase.org).

## Results

### Phenotypic variation of early and late leaf spot diseases

For ELS, GP-NC WS 16 had lower disease scores than Florida-07 in both 2013 (P<0.05) and 2014 (P=0.08) (Table S1). For LLS, the disease rating for GP-NC WS 16 was significantly lower than Florida-07 in both 2012 and 2015 (P<0.05). No statistical difference between the parents was found in 2013 and 2014. As for yield data collected from the LLS field trials, no statistical difference was found between the two parental lines across all three years. The GA subpopulation showed near to normal distribution for both diseases and yield (Figure 2). Analysis of variance test for ELS showed significant difference among RIL (F=12.5, P<0.0001), environment (F=2768.2, P<0.0001) and RIL x environment interaction (F=1.4, P=0.0048). The broad sense heritability for ELS was 0.4. As for LLS, significant difference was found among RIL (F=7.7, P<0.0001) and environment (F=57.7, P<0.0001) whereas the RIL x environment interaction (F=1.0, P=0.34) was not significant. The broad sense heritability for LLS was 0.8. Analysis of variance test for yield showed significant difference among RIL (F=4.9; P<0.0001), environment (F=870.9, P<0.0001) and RIL x environment interaction (F=1.7; P<0.0001). The broad sense heritability for yield was 0.6. Correlation analysis indicated that ELS disease ratings for 2013 and 2014 were correlated at 0.66 (p<0.01). Since the yield data were collected following the LLS disease ratings, correlation among the four years of LLS disease scores, three years of yield and their respective combined data was tested (Table 1). Highly significant positive correlation among the LLS disease ratings was found ranging from 0.48 to 0.90 (p<0.01). Within the three year yield data, significant positive correlation ranging from 0.32 to 0.86 was found. More importantly, highly significant negative correlation (−0.17 to −0.52) was detected among most of the yield and LLS disease ratings except for correlation among yield 2013, LLS 2013 and LLS 2014 ratings which did not reach statistical significance.

**Table 1.**
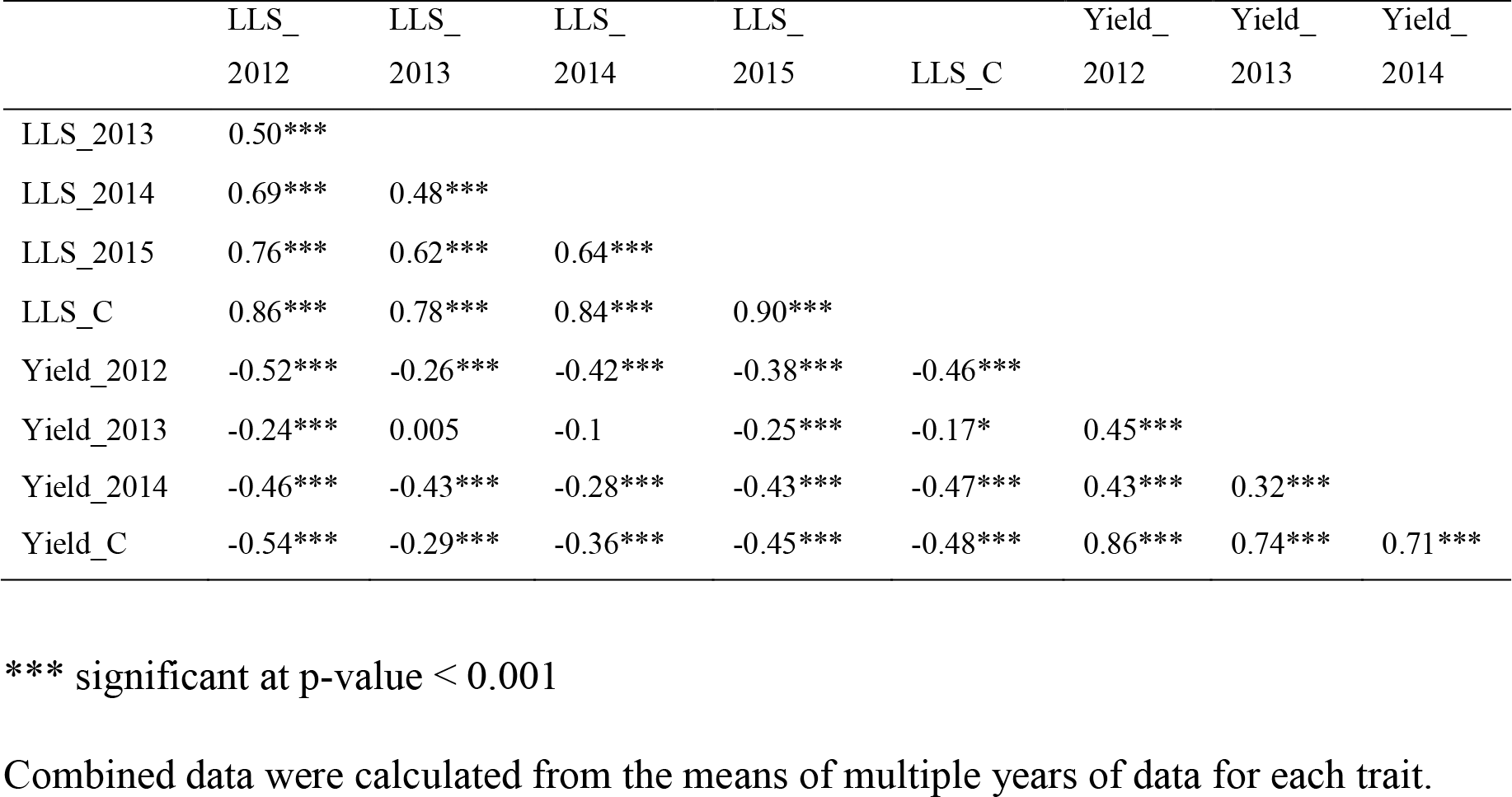
Correlation of late leaf spot (LLS), combined late leaf spot (LLS_C) disease ratings, yield, and combined yield (yield_C) collected from the Georgia subpopulation in the non-sprayed field tests.

**Table 2.**
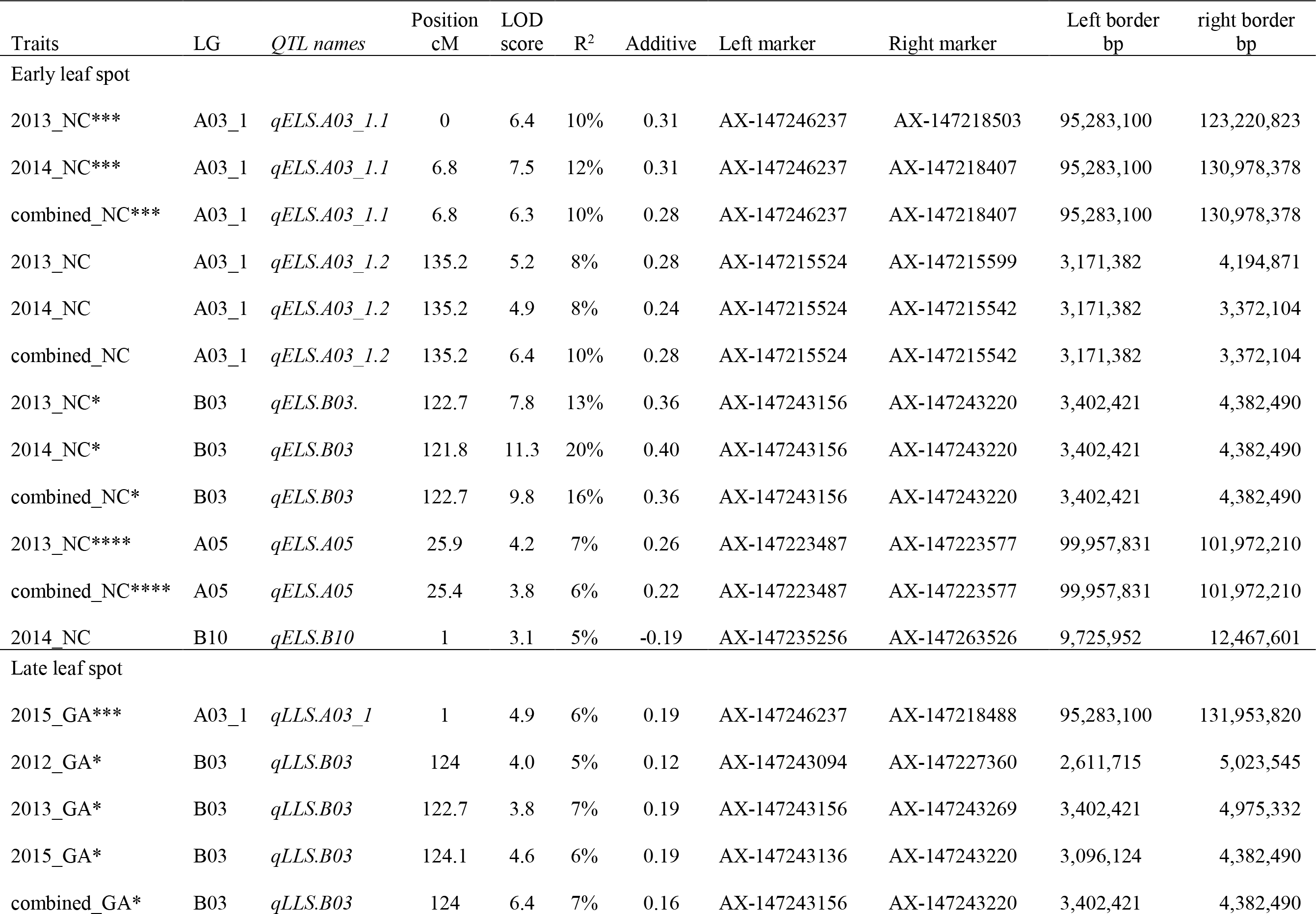
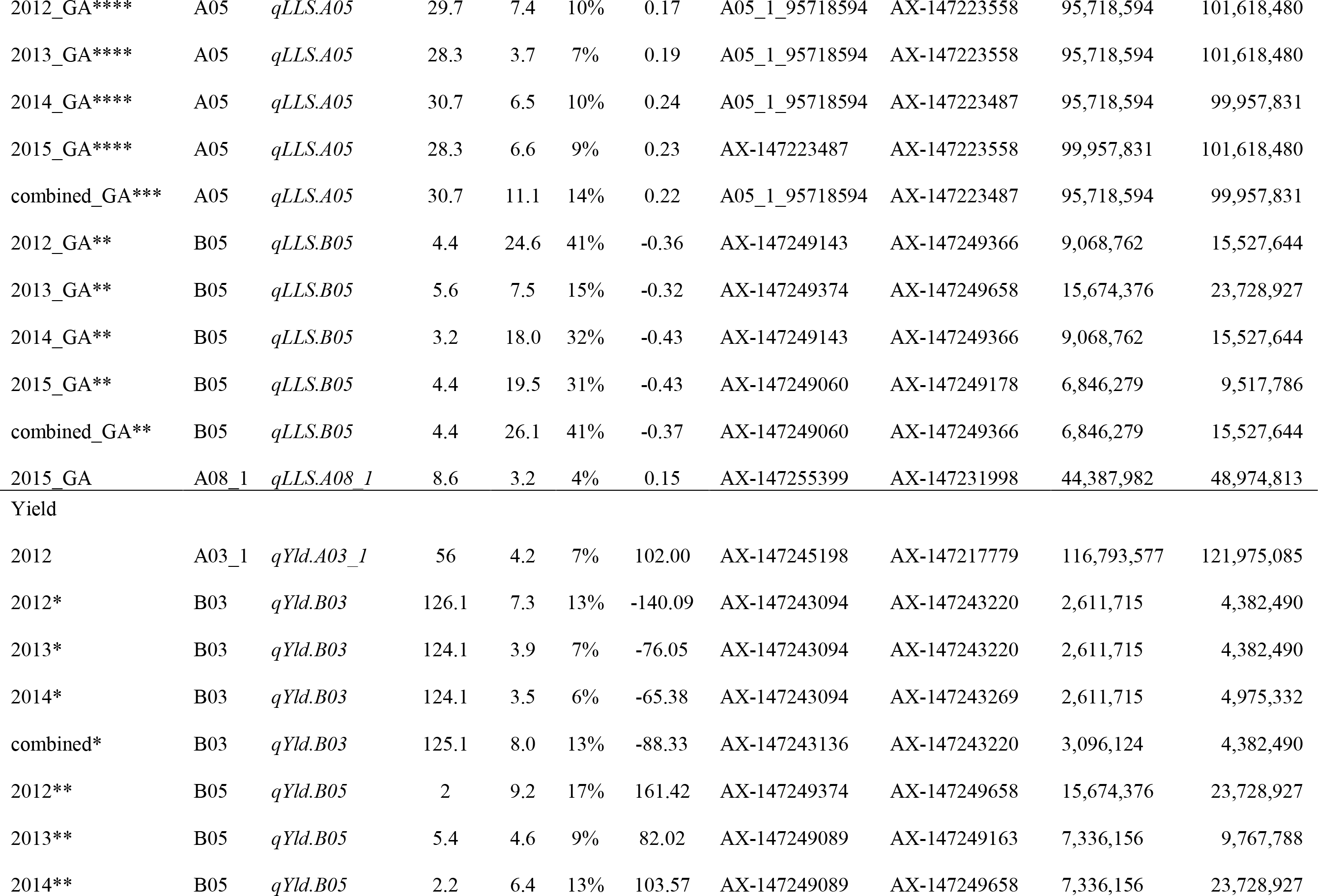
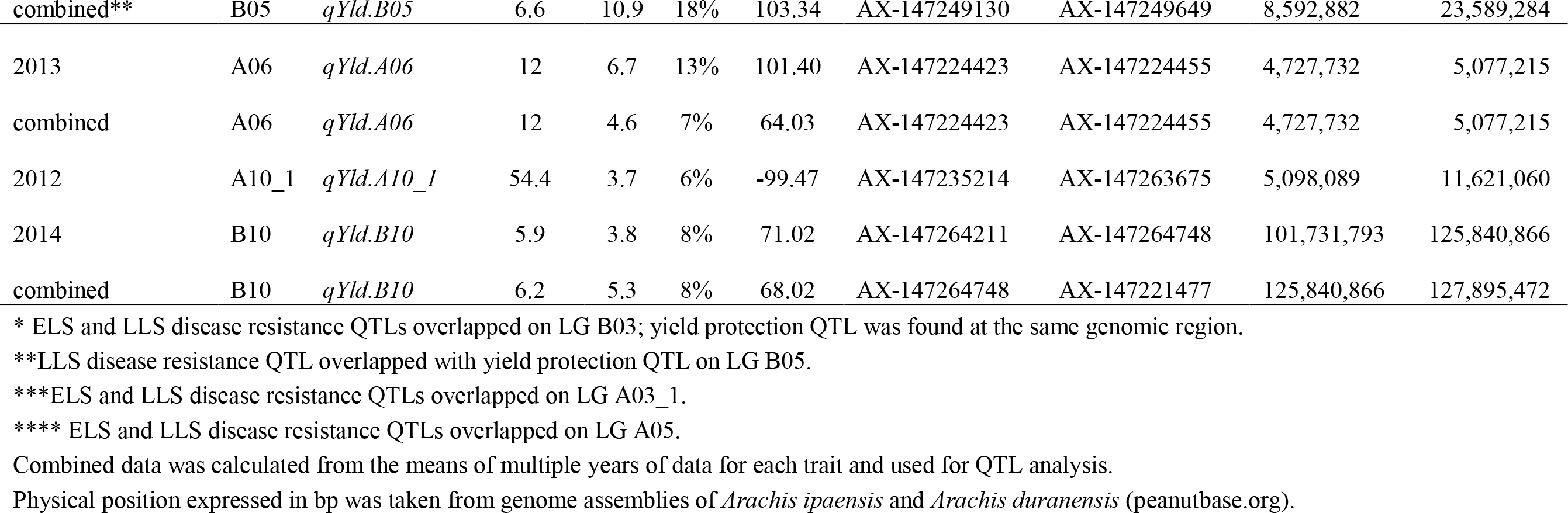
QTLs detected for early and late leaf spot disease resistance and their impact on yield using the Georgia subpopulation.

**Figure 2.**
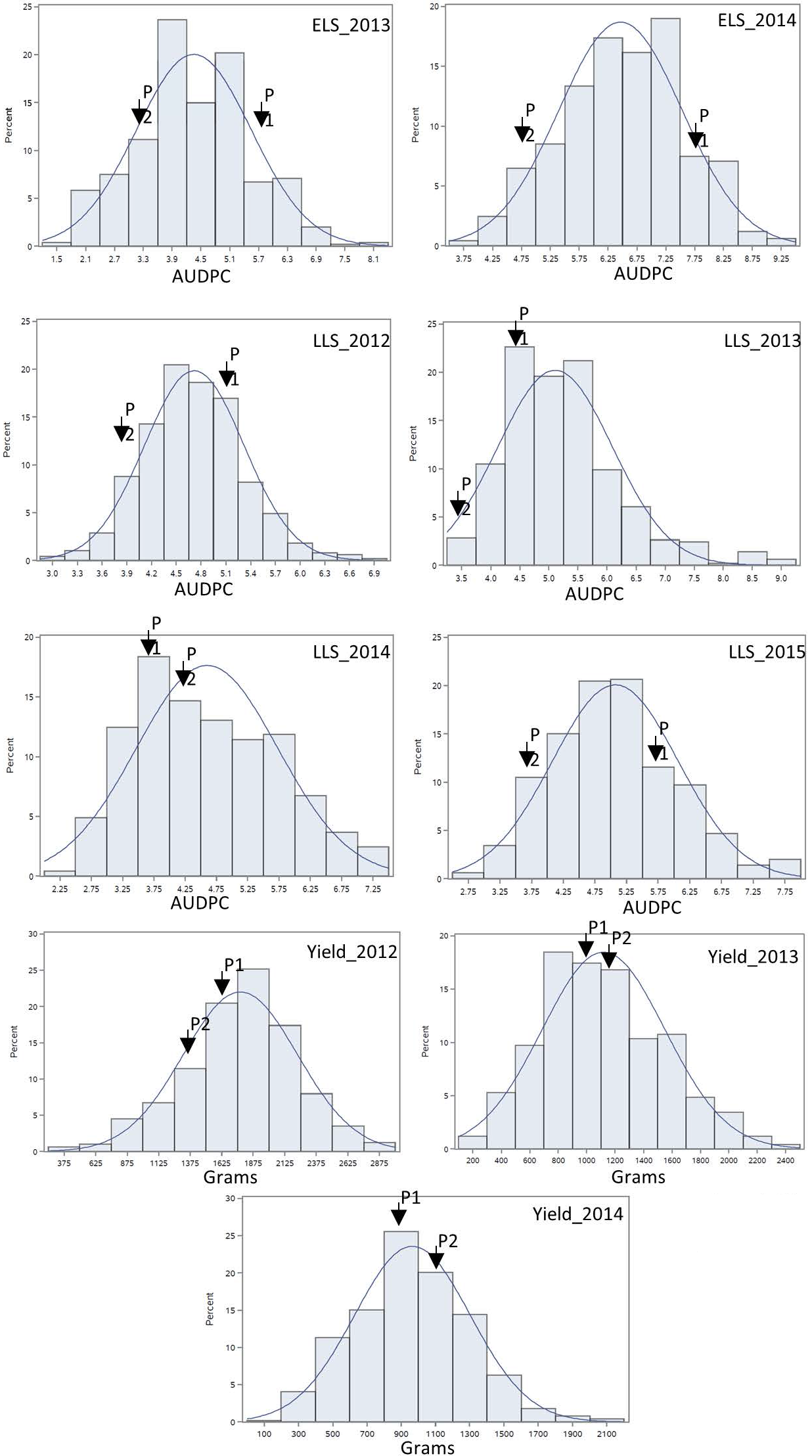
Phenotypic distribution of area under the disease progress curve of ELS and LLS diseases and yield collected in non-sprayed test fields. P1: female parent Florida-07; P2: male parent GP-NC WS 16. The normal distribution curve represents the expected distribution of disease score and yield for the RIL population.

### Genetic markers and linkage map construction

Ninety-one percent of the 58,233 loci in the Axiom SNP array were monomorphic between the GP-NC WS 16 and Florida-07 parent and 999 SNP markers were curated for this population. Sixty-three SSR loci were used for genetic mapping (Table S2), of which 58 loci had been previously mapped (http://marker.kazusa.or.jp/Peanut) and all but 7 loci mapped to the expected LG assignment in our map. The SSRs were distributed across most chromosomes except for A04, B07 and B09. The high oleic marker ahFAD2B_hybprobe was mapped to LG B09 as expected. In addition, four of the six KASPar markers were also mapped to the expected LGs while the other two mapped to their respective homeologous LGs. Consequently, a genetic map consisting of 855 loci distributed on 28 linkage groups (LGs) covering 1414.8 cM was constructed (Table S3, Figure S1). Fifteen LGs were assigned to the A subgenome and 13 LGs to the B subgenome based on the physical positions of the SNP markers. The average map density was 2.2 cM/locus ranging from 0.3 to 8 cM/locus whereas the largest gap between adjacent loci was 23 cM on LG A03_1. Marker distortion was not detected on most LGs except in LG A10_1 with 71% of loci distorted towards GP-NC WS 16 allele.

In order to detect if there is any wild introgression from *A. cardenasii* retained in GP-NC WS16, the Intromap SNP calling pipeline was used to discover diagnostic SNPs for introgression. The results failed to show any significant signals indicating the wild segment is either lost in the process of material advancement or the segments were reduced to an undetectable size using the Intropmap SNP calling pipeline.

### QTL mapping for ELS, LLS and Yield

A total of five QTLs were detected for ELS disease resistance with three QTLs mapped to the A subgenome and two to the B subgenome (Table 2, Figure 3). The QTL *qELS.A03_1.1* (phenotypic variance explained, PVE=10 to 12%), *qELS.A03_1.2* (PVE=8 to 10%), and *qELS.B03* (PVE=13-20%) were detected in both 2013 and 2014 single year data as well as in the combined datasets. These QTLs are considered consistent major QTLs. The QTL *qELS.A05* (PVE=6-7%) was detected in 2013 and combined datasets whereas *qELS.B10* (PVE=5%) was detected only from the 2014 disease data. The resistance alleles for all five QTLs came from the GP-NC WS 16 parent.

**Figure 3.**
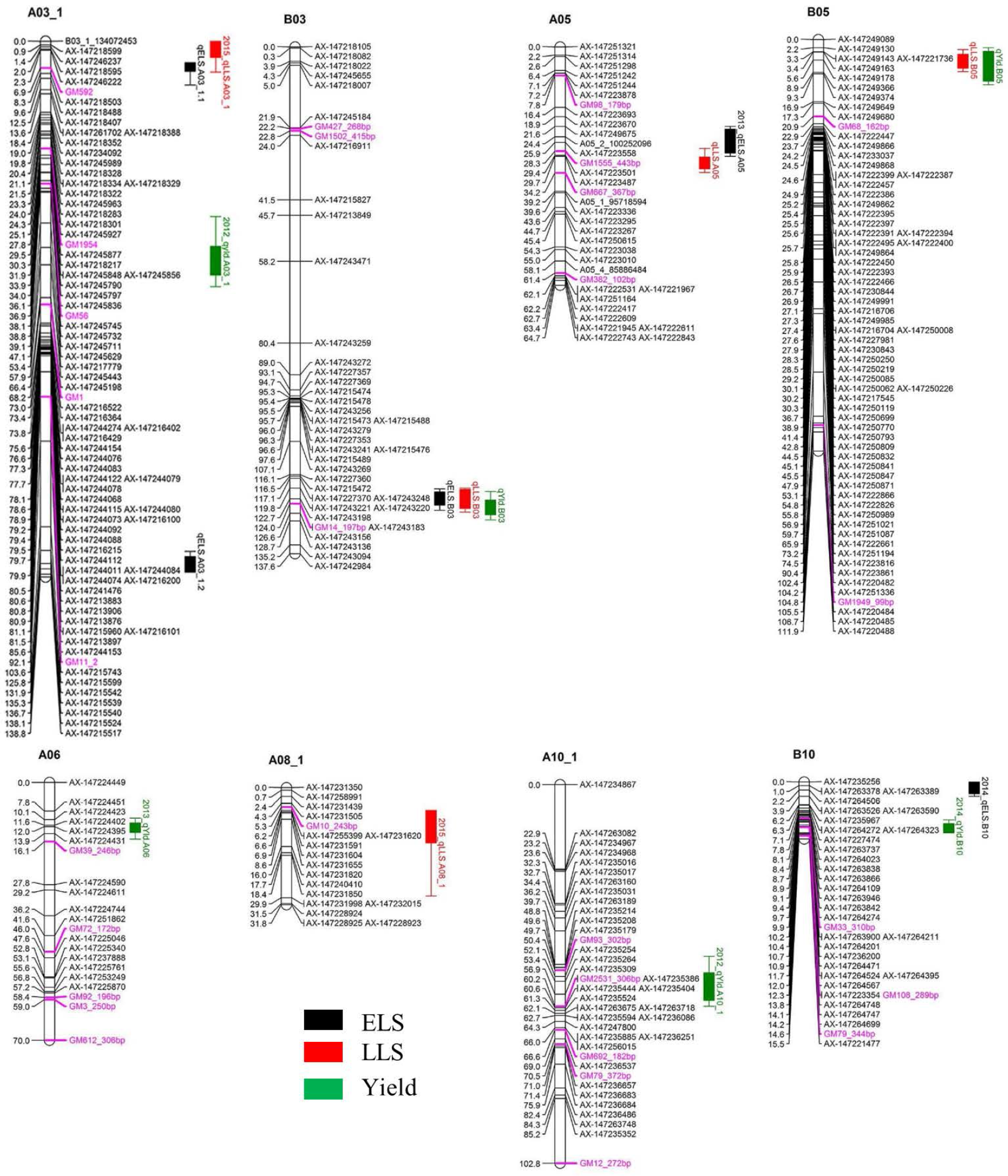
QTLs for early leaf spot (ELS), late leaf spot (LLS) disease resistance and yield detected in the Georgia-advanced RIL subpopulation. Map distance (cM) is shown to the left of each linkage group and genetic markers on the right. Traits without year suffix were identified by combined data analysis. Single year QTLs that overlapped with QTLs identified by combined data are not shown on the graph. SSR markers are indicated by purple font color.

Five QTLs also were detected for LLS disease resistance, three on the A subgenome and two on the B subgenome. The QTL *qLLSB03* (PVE=5-7%) was detected in 2012, 2013, and 2015 single year data and the combined datasets. The QTL *qLLSA05* (PVE=7-10%) and *qLLSB05* (PVE=15-41%) were identified in all four single year data and combined datasets.

These three QTLs were considered consistent QTLs. The QTL *qLLSA03_1* (PVE=6%) and *qLLSA08_1* (PVE=4%) were detected in the 2015 dataset only. Except for *qLLSB05* whose resistant allele came from Florida-07, the remaining four QTLs had GP-NC WS 16 contributing the resistant allele.

Two genomic regions showed significant association with both ELS and LLS resistance (Table 2, Figure 3). In linkage group A03, the locus *qELSA03_1.1* was mapped to the region that overlapped with *qLLSA03_1*, which was flanked by the markers AX-147246237 and AX-147218488. Similarly, the genomic region of *qELSA05* overlapped with *qLLSA05* on linkage group A05 near the flanking markers AX-147223487 and AX-147223558. For all QTL regions, the parental allele which provided resistance to ELS also contributed resistance to LLS.

Six QTLs were detected for yield with equal distribution on the A and B subgenomes. The QTL *qYldB03* (PVE=7%) and *qYldB05* (PVE=9-13%) were detected in all three single year and combined datasets, therefore they were considered consistent QTLs. The QTL *qYldA03_1* (PVE=7%) and *qYldA10_1.1* (PVE=6%) were detected only in 2012 data and *qYldA06* (PVE=13%) and *qYldB10* (PVE=8%) were detected in 2013 and 2014 data, respectively, as well as in the combined dataset. For *qYld.B03* and *qYld.A10_1*, the favorable allele came from the GP-NC WS 16 parent while for the other four QTLs, the Florida-07 allele improved yield.

### Correspondence of disease resistance and yield QTLs

Several QTL regions for resistance to ELS and/or LLS corresponded to QTL for yield (Table 2, Figure 3). For example, the disease resistance loci *qLLS.B03* and *qELSB03* were mapped to the same genomic region as the yield locus *qYldB03* on linkage group B03 flanked by the markers AX-147243156 and AX-147243220. Also on linkage group B05, the LLS resistance locus *qLLSB05*, which was consistently detected in all single years and the combined dataset and explained up to 41% of PVE, was mapped to the same region as *qYldB05*, the most significant yield QTL that explained up to 18% PVE. Finally, the resistance locus *qELS.B10* was mapped to the genomic region that overlapped with the yield locus *qYldB10* on linkage group B10. For all QTL regions, the parental allele in which provided resistance also contributed positively to yield.

### Confirmation of ELS resistance QTLs

The NC subpopulation was utilized in a post hoc analysis to determine the effectiveness of screening with flanking markers for the three consistent ELS resistance QTLs (*qELS.A03_1.1*, *qELS.A03_1.2* and *qELS*.B03) to select for resistant and susceptible genotypes based on the 2015 data collected in North Carolina. The mean disease rating of the genotypic classes with and without the resistant allele from the GP-NC WS 16 parent for each QTL is shown in Figure 4. For *qELS.A03_1.2* and *qELS.B03*, the mean rating of the RILs carrying the resistant allele was significantly lower than those carrying the susceptible allele. While no significant difference was found between RILs grouped by markers flanking *qELS.A03_1.1,* the RILs with the resistant allele were numerically lower than those with the susceptible allele. For combined data across the three QTLs, the RILs possessing resistant GP-NC WS 16 alleles clearly had a significantly lower ELS disease rating than those with the susceptible Florida-07 alleles.

**Figure 4.**
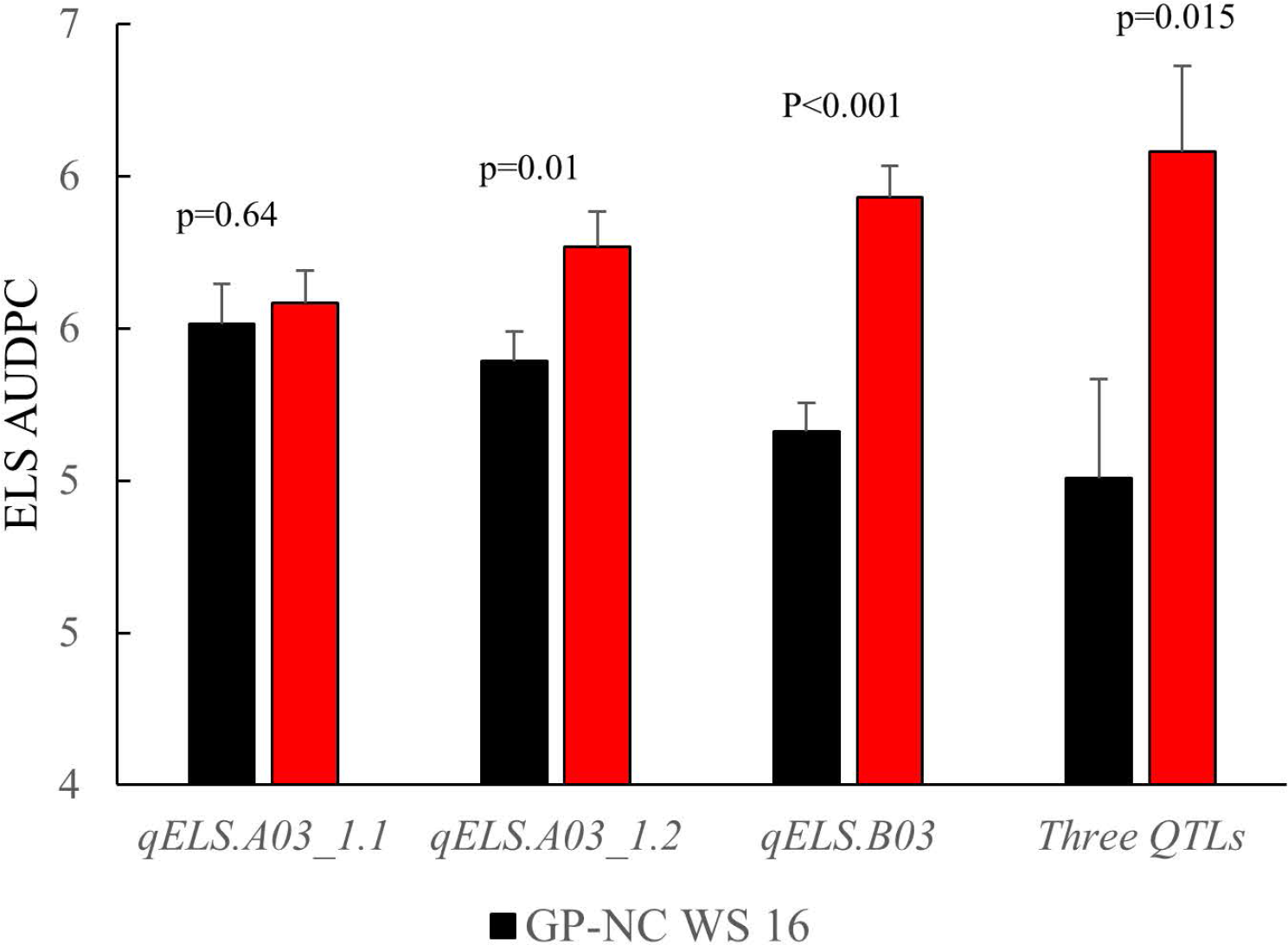
Confirmation of QTLs identified for early leaf spot disease. Flanking markers of early leaf spot QTLs were used to select RILs from the NC subpopulation to produce to two groups, i.e., homozygous for Flordia-07 or GP-NC WS 16 alleles, respectively. Year 2015 early leaf spot disease data from these two groups of RILs were compared by t-test according to their parental allele profile. From left to right, each set of RILs was selected by one of three consistent QTLs, i.e., *qELS.A03_1.1, qELS.A03_1.2* or *qELS.B03,* respectively. The last set of RILs was selected by all three QTLs.

### Gene content of consistent QTL regions

By retrieving diploid genome gene models across physical intervals of the five consistent QTLs, a total of 2,184 gene models were compiled, 69 of which are annotated with a potential function in disease resistance (Table S4). Within the genomic region of *qELS.A03_1.1*, there were 1,042 gene models, 36 of them in the families of lipoxygenase, serine threonine-protein phosphatase and WRKY family transcription factor. As for *qELS.A03_1.2*, there were 120 gene models, and CC-NBS-LRR disease resistance protein and serine threonine-protein phosphatase were among them. The overlapping QTL region of *qELS.B03* and *qLLS.B03* contained 264 gene models. Potential disease resistance genes in this region include dirigent-like disease responsive protein and serine threonine-protein phosphatase. There were 270 and 487 gene models within the QTL regions of *qLLS.A05 and qLLS.B05*, respectively. Serine threonine-protein phosphatase is a potential candidate for disease resistance in these regions.

## Discussion

Although early and late leaf spot diseases occur naturally in peanut growing regions, the disease epidemic and severity is influenced by the history of leaf spot incidence, crop rotation and fungicide application (Fulmer, 2017). Both early and late leaf spot data sets collected for the GA subpopulation demonstrated significant environmental effects suggesting the disease pressure varied across multiple years of field tests conducted in Georgia. For LLS, the GA subpopulation responded consistently across the four-year tests since no significant RIL x environment interaction was found, whereas significant RIL x environment interaction was found for ELS disease ratings suggesting that the disease ratings of the RIL were influenced by the year of data collection. Regardless, significant positive correlations were found for disease ratings between different years for each leaf spot disease indicating consistent disease response of RILs across environments. Based on the complicity of factors impacting the phenotyping data, QTL mapping was performed using the individual year datasets separately and the combined across years dataset.

The power to identify QTLs can be improved with higher density genetic maps, increased population size, and more accurate collection of phenotypic data (Broman et al., 2001). Earlier QTL mapping experiments by using the SSR-based maps have low marker density ranging from only 56 to 418 loci (Khedikar et al., 2010; Sujay et al., 2012; Wang et al., 2013; Khera et al., 2016; Pandey et al., 2017b). Recently, two SNP-based maps with more than 1,000 loci have been developed to map QTLs for resistance to LLS and ELS diseases (Zhou et al., 2016; Liang et al., 2017); however, the physical positions of SNP markers were not reported rendering no basis for comparison with current work. Our linkage map placed 855 marker loci with 92% of the loci being new SNPs markers from the publicly available Axiom_*Arachis* 58K SNP array and the rest from previously mapped SSRs, KASPar, and the functional marker for *ahFAD2B*. This new linkage map provides markers for both leaf spot diseases that can be used in breeding programs using related sources of resistance. In addition, the use of AUDPC data as well as four years of phenotyping have provided highly robust datasets for the LLS disease compared to most other mapping studies.

The number of resistant loci reported in previously published studies for leaf spot disease ranged from 6 to 37 QTLs. While the genetic populations used in these studies may harbor more alleles contributing to leaf spot disease resistance, a majority of the QTLs have not been validated. Among the five QTLs identified for LLS, *qLLS.B03*, *qLLS.A05* and *qLLS.B05* were consistently detected by multiple years of disease ratings. Further, QTL regions, *qLLS.A05* and *qLLS.B05,* confirmed our previous results using QTL-seq analysis (Clevenger et al., 2018). The locus *qLLS.B03* covers a 2.6 to 5.0 Mbp genomic region on chromosome B03 which was not identified previously by QTL-seq. Finally, the locus *qLLS.A03_1* (95 to 132 Mbp) detected only from the 2015 data overlapped with the B03 LLS resistant region reported by QTL-seq analysis. Peanut is an allotetraploid with 96% of sequence similarity between the two subgenomes (Bertioli et al., 2016). Further study will be needed to resolve the discrepancy of subgenome assignment. The last minor QTL *qLLS.A08_1* was only found by genetic mapping. QTL-seq is an extension of bulk segregant analysis which detects the common SNPs between the resistant and susceptible bulks based on sequencing analysis. Confirmation of the two consistent QTLs between these two methods reassures the validity of our QTLs. Although QTL-seq is more rapid in QTL identification, its estimation of QTL effect is not as thorough as composite interval mapping since it only includes a subset of the population for sequence analysis.

As for ELS, out of a total of five QTLs identified, three were detected across multiple environments supporting the robustness of the marker-trait association for these genomic regions (Table 2, Figure 3). The post hoc analysis with the NC subpopulation demonstrated the effectiveness of these QTL regions in selecting for resistant lines in breeding populations (Figure 4). The disease score dropped by an average of 0.33 points between the RILs harboring the resistant allele compared to the lines with the susceptible allele selected by individual QTLs. When the RILs from the NC subpopulation were selected with all flanking markers of all three QTLs, the largest phenotypic difference was achieved - the disease dropped 1.63 points with lines harboring resistant alleles - suggesting the positive additive effect of these QTLs. While the flanking markers used for genotype selection are SNP array markers, they can easily be converted to the high throughput KASPar assays to provide a smooth transition for utilization these markers in breeding programs (Chu et al., 2016; Clevenger et al., 2018).

As expected, the GP-NC WS 16 parent, an interspecific introgression line developed for ELS resistance at NCSU (Stalker, 2017), was the source of resistance alleles for all of the ELS QTLs. Therefore, while we cannot definitively ascribe any of the GP-NC WS 16 genome to wild introgression from *A. cardenasii*, it is quite possible that the source of one or more of the ELS resistance alleles we identified could come from this diploid relative. Interestingly, locus *qELSA03_1.1* (123 to 131 Mbp) and *qLLSA03_1* (95 to 132 Mbp) corresponds with a QTL previously mapped in the LLS resistant line GPBD 4 (Pandey et al., 2017a), which was derived from the same wild introgression line as GP-NC WS 16. However, unlike GPBD 4, GP-NC WS 16 had lost most of its introgression from *A. cardenasii* due to many generations of backcrossing while under field selection for early leaf spot resistance, resulting in likely retention of only a small portion of alien chromatin on chromosome A03 conferring ELS resistance (Tallury et al., 2014).

Both ELS and LLS agents have the ability to over winters in the soil where the conidia are deposited on the debris of plant tissue. Beginning around mid-season, the pathogens progressively encroach upon peanut plants starting from the leaves closest to the ground and migrating to the upper layers of the canopy. If fungicides are not applied after the appearance of symptoms, both ELS and LLS will cause defoliation towards the later stages of disease progression. Although disease ratings were recorded at two separate locations with LLS being the predominant disease in the Georgia location and ELS in the North Carolina location, the co-existence of both diseases has been observed in both states. Therefore, correspondence of QTLs between the two diseases is not unexpected in this study. Indeed, three QTL regions were found to overlap between the two disease resistance traits: *qELS.A03_1.1* and *qLLSA03_1*; *qELS.B03* and *qLLS.B03*; *qELSA05* and *qLLSA05*. Because the pathogens of both early and late leaf spot are host specific, infecting only the genus *Arachis*, and share a similar life cycle, it is possible that similar genetic mechanisms of host resistance could provide protection against both fungal diseases. Alternatively, these co-located resistance QTLs for ELS and LLS could be due to genetic linkage.

When the condition is favorable for infection, leaf spot lesions begin to appear within 3-5 weeks after planting for ELS and about one month later for LLS. Since it only takes 10 to 15 days for the newly emerged lesions to sporulate, both diseases can go through many cycles of reproduction before harvest; therefore, with no fungicide applications, both leaf spot diseases can result in severe pod yield loss. In this study, the field evaluation was conducted without fungicide applications, which created environmental conditions that were highly favorable to disease development. The negative correlation between yield and LLS disease rating in the four years of evaluation in Georgia suggests that LLS disease played a major role in yield reduction (Table 1). Interestingly, two yield QTLs *qYld.B03* and *qYld.B05* corresponded with the major QTLs for LLS resistance *qLLS.B03* and *qLLS.B05*, respectively. In addition, for both QTL regions, the alleles from GP-NC WS 16 and Florida-07 provided resistance to LLS as well as yield improvement, indicating that the resistance alleles have protected against yield losses caused by damage due to the LLS disease. While it is also possible that bonafide yield QTLs could be present in these chromosome regions (Huang et al., 2015; Chen et al., 2017; Luo et al., 2017), direct comparisons of QTL positions were not possible because no physical position was provided in the previous studies. Future field trials with and without fungicide management will help clarify if the yield QTLs identified herein were responding to or independent of the resistance alleles for LLS disease.

The genomic regions defined by the consistent QTLs range from 1 to 38 Mbp and contain over 2,000 gene models. Several disease resistance genes within these regions may have an impact on the host response to leaf spot infection. Serine threonine-protein phosphatase has been shown to negatively regulate plant defense against fungal infection (Pais et al., 2009). Suppression of its activity is required to activate the localized cell death response (He et al., 2004). There are 59 gene models of this protease within the *qELS.A03, qLLS.A05* and *qLLS.B05,* often in tandem arrays along the chromosomes. Tandem and segmental duplications of plant disease resistance genes has been reported previously in *Arabidopsis* (Leister, 2004). Although large in copy number, only two gene models of this protease within *qELS.A03* and *qLLS.B05* demonstrated gene expression in the Tifrunner transcriptome database (Clevenger et al., 2016; peanutbase.org) suggesting that most may be pseudogenes. WRKY family transcription factor was suggested to play a central role in activating plant defense systems (Eulgem and Somssich, 2007). Four gene models of WRKY transcription factors highly expressed in peanut were found in the *qELS.A03* genomic region. A lipoxygenase was found within *qELS.A03* and lipoxygenases are also known play a role in plant defense by producing phyto-oxylipins (Blee, 2002). There were three dirigent-like protein genes on *qELS.B03*. These proteins were reported to be involved in induced phenolic plant defense mechanisms (Ralph et al., 2006). Further studies will be needed to determine the relevance of these defense genes to leaf spot resistance.

In summary, with the aid of a SNP-based genetic map, consistent QTLs for resistance to ELS and LLS diseases were identified on chromosomes 3 and 5, respectively. QTLs for ELS were independently confirmed by the power of correct phenotypic separation of resistant and susceptible RILs defined by flanking markers using a subpopulation not involved in marker-trait association analysis. The locus *qELSA03_1.1* and *qLLS.A03_1* corresponded with the late leaf spot resistant region identified previously in the resistant line GPBD 4. As for LLS QTLs, the genetic mapping results were consistent with the published QTL-seq analysis. These resistance QTLs may have protected the yield loss caused by late leaf spot disease. Implementing marker-assisted selection holds promise for pyramiding of both ELS and LLS resistance into current elite peanut cultivars.

## Author Contributions

PO-A, YC, TI and CC designed the experiments; TI and CC provided the genetic materials; YC, AC, TI performed the disease ratings; YC and PC wrote the original draft and were responsible for data visualization; PO-A administer the project.

## Acknowledgement

Xuelin Luo provided statistical analysis and Josh Clevenger performed IntroMap analysis. Technical support was received from Shannon Atkinson, Stephanie Botton, Jason Golden, Kathy Marchant, Brant Sandifer, Betty Tyler and Congling Wu. The authors gratefully acknowledge their dedicated contribution to this project. This work was supported in part by funds from The Peanut Foundation, the Georgia Peanut Commission, and The University of Georgia Research Foundation Cultivar Development Research Program.

