## Supplemental Figure 1 for "Major QTLs for resistance to early and late leaf spot diseases are identified on chromosomes 3 and 5 in peanut (*Arachis hypogaea*)"

### Slide 1
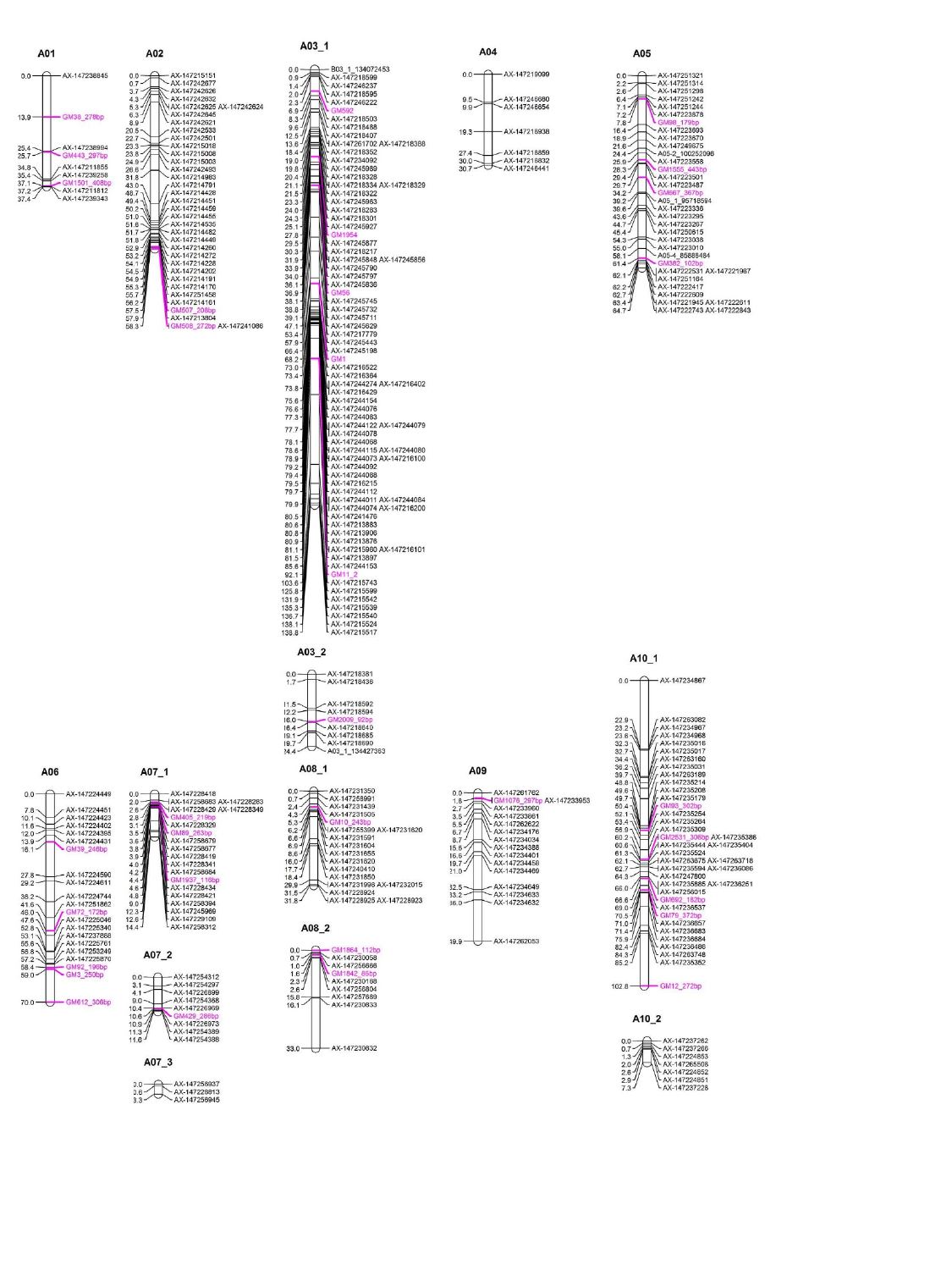

### Slide 2
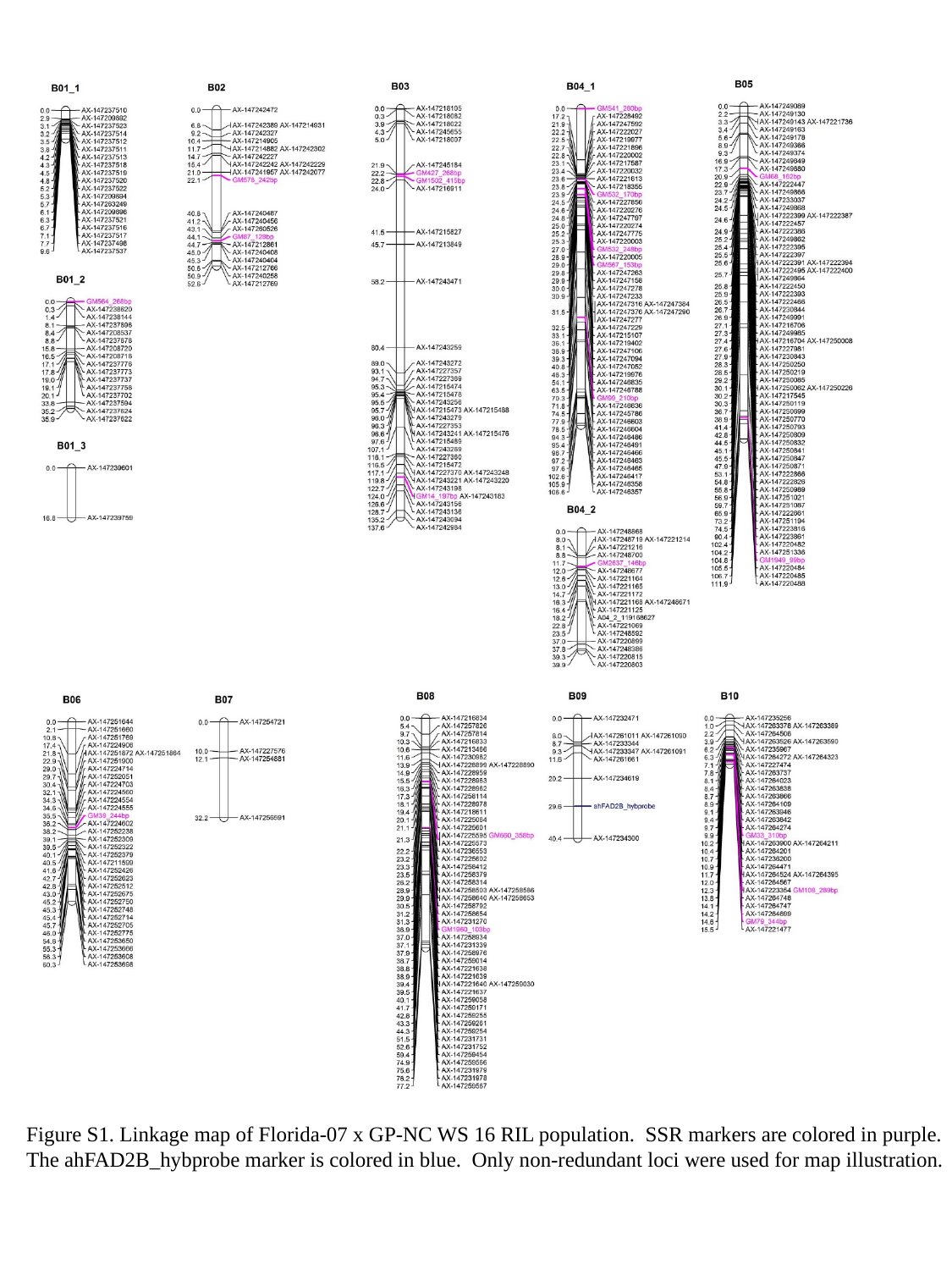

Figure S1. Linkage map of Florida-07 x GP-NC WS 16 RIL population. SSR markers are colored in purple.
The ahFAD2B_hybprobe marker is colored in blue. Only non-redundant loci were used for map illustration.
