## Supplemental Tables 1-3 for "Major QTLs for resistance to early and late leaf spot diseases are identified on chromosomes 3 and 5 in peanut (*Arachis hypogaea*)"

| Table S1. Disease data summary of the Georgia subpopulation. | | | | | | | | | | | |
| --- | --- | --- | --- | --- | --- | --- | --- | --- | --- | --- | --- |
| **traits** | **Florida 07** | **GP-NC WS 16** | **RIL Range** | **Mean** | **SD** | **Skew** | **Kurt** | | **Shapiro-Wilk (W)** | | **Pr < W** |
| ELS_2013 | 5.7 | 3.3 | 2.1-7.2 | 4.4 | 1.2 | 0.16 | -0.35 | | 0.99 | | 0.0039 |
| ELS_2014 | 7.4 | 5.1 | 4.6-8.6 | 6.5 | 1.1 | -0.10 | -0.41 | | 0.99 | | 0.0105 |
| LLS_2012 | 4.9 | 3.8 | 2.6-6 | 4.7 | 0.6 | 0.29 | 0.44 | | 0.99 | | 0.0263 |
| LLS_2013 | 4.8 | 3.6 | 3.6-7.3 | 5.1 | 1.0 | 1.06 | 1.62 | | 0.93 | | <0.0001 |
| LLS_2014 | 3.6 | 4.2 | 2.1-5.7 | 4.6 | 1.1 | 0.35 | -0.70 | | 0.97 | | <0.0001 |
| LLS_2015 | 5.8 | 3.9 | 3.5-7.5 | 5.1 | 1.0 | 0.39 | -0.03 | | 0.99 | | 0.0002 |
| Yield_2012 | 1702 | 1442 | 709.3-2726.7 | 1773.0 | 453.4 | -0.28 | 0.12 | | 0.99 | | 0.0192 |
| Yield_2013 | 1014 | 1152 | 578.7-1863.3 | 1118.8 | 431.9 | 0.33 | -0.34 | | 0.99 | | 0.0003 |
| Yield_2014 | 965 | 1201 | 298-1756 | 965.4 | 338.5 | 0.25 | 0.04 | | 0.99 | | 0.0289 |
| ELS stands for early leaf spot disease ratings; LLS stands for late leaf spot disease ratings. | | | | | | | | | | |  |
| Disease ratings were expressed as area under disease progression curve (AUDPC). | | | | | | | | |  | |  |
| Yield was measured as gram/plot. | | | | | | | | | | | |

| Table S2. Comparison of SSR marker linkage group assignment between current map | | | |
| --- | --- | --- | --- |
| and previously published maps for *Arachis hypogaea*. | | | |
| Published linkage group assignment information collected from http://marker.kazusa.or.jp/Peanut | | | |
| Linage group | SSR markers | SSR alias | Linage group assignment from published maps* |
| A03_1 | GM1_285bp | TC0A01 | A03 |
| A08_1 | GM10_243bp | TC1E05 | A08 |
| A09 | GM1076_297bp |  | A07; B07 |
| B10 | GM108_289bp | AC2B03 | B10; A10 |
| A03_1 | GM11_141bp | TC1E06 | B03; A03 |
| A03_1 | GM11_151bp | TC1E06 | B03; A03 |
| A10_1 | GM12_272bp | TC1G04 | A10; B10; B05 |
| A02 | GM126_389bp | RI1F06 | A02; B02 |
| B03 | GM14_197bp | TC2A021 | B03 |
| A01 | GM1501_408bp |  | B01 |
| B03 | GM1502_415bp |  | B03 |
| A05 | GM1555_443bp |  | B05 |
| B05 | GM1577_288bp |  | A05 |
| A08_2 | GM1842_86bp |  |  |
| A08_2 | GM1864_112bp |  |  |
| A07_1 | GM1937_116bp |  | A07 |
| B05 | GM1949_99bp |  |  |
| A03_1 | GM1954_125bp |  | A03; B03 |
| B08 | GM1960_103bp |  |  |
| A03_2 | GM2009_92bp |  | B03 |
| A10_1 | GM2531_306bp |  | A10; B10 |
| B04_1 | GM2589_335bp |  | A04; B04 |
| B04_2 | GM2637_146bp |  | A04 |
| A06 | GM3_250bp | TC1A02 | A06; B03; B06; B07 |
| B10 | GM33_310bp | TC3E05 | B10 |
| A01 | GM38_278bp | TC3H02 | A01 |
| A05 | GM382_102bp | PM45 | A05 |
| B06 | GM39_244bp | TC3H07 | A06; B06 |
| A06 | GM39_246bp | TC3H07 | A06; B06 |
| A07_1 | GM405_219bp | PM204 | A07 |
| A03_1 | GM415_146bp | PM238 | B03; A03 |
| B08 | GM423_254bp | Seq2A06 | B08; A07 |
| B03 | GM427_268bp | Seq2C11 | B03; A03 |
| A07_2 | GM429_286bp | Seq2E06 | A07; B03; B07 |
| A01 | GM443_297bp | Seq3B05 | A01 |
| B10 | GM450_275bp | Seq3E10 | B10; A10 |
| A02 | GM507_208bp | Seq11G03 | A02 |
| A02 | GM508_272bp | Seq11G07 | A02; B08 |
| B04_1 | GM532_170bp | Seq14D11 | B04; B02; B07 |
| B04_1 | GM532_248bp | Seq14D11 | B04; B02; B07 |
| B04_1 | GM541_260bp | Seq15C12 | B04; A04 |
| A03_1 | GM56_180bp | TC4G02 | B03; A03 |
| B01_2 | GM564_268bp | Seq17E01 | B01 |
| B04_1 | GM567_153bp | Seq17F06 | B04 |
| B02 | GM578_242bp | Seq18E07 | B02 |
| A03_1 | GM592_297bp | Seq19D09 | B03; A03; A09 |
| A06 | GM612_285bp | Seq9H08 | A03; B06 |
| A03_1 | GM612_306bp | Seq9H08 | A06; A03 |
| A10_1 | GM647_285bp |  | A10 |
| B05 | GM657_350bp |  | B05 |
| B08 | GM660_358bp |  | B08 |
| A05 | GM667_367bp |  | A05 |
| B05 | GM68_162bp | TC6E01 | A05; B05 |
| A10_1 | GM692_182bp |  | A10 |
| A06 | GM72_172bp | TC7C06 | A06; B10 |
| B10 | GM79_344bp | TC7H11 | B10; A10 |
| A10_1 | GM79_372bp |  |  |
| B02 | GM87_128bp | TC9F04 | B02; A03; A08 |
| A07_1 | GM89_263bp | TC9H08 | A07 |
| A06 | GM92_196bp | TC11A04 | A06; B06; A10 |
| A10_1 | GM93_302bp | TC11B04 | A10; B05; A04 |
| A05 | GM98_179bp | TC11F12 | A03; A08; B07 |
| B04_1 | GM99_210bp | TC11A06 | B04 |

| Table S3. Description of the linkage map of Florida 07 x GP-NC WS 16 recombinant inbred population | | | | | | | | | |
| --- | --- | --- | --- | --- | --- | --- | --- | --- | --- |
|  |  |  |  |  | segregation distorted loci (SDL) | | | |  |
| LG name | map distance (cM) | loci number All | Largest gap (cM) | map density (cM/loci) | SDL | SDL% | Female parent | Male parent | loci number non-redundant |
| A01 | 37.4 | 10 | 13.9 | 3.7 | 0 | 0.0 | 0 | 0 | 9 |
| A02 | 58.3 | 40 | 11.6 | 1.5 | 0 | 0.0 | 0 | 0 | 35 |
| A03_1 | 138.6 | 98 | 23.1 | 1.4 | 0 | 0.0 | 0 | 0 | 79 |
| A03_2 | 24.4 | 9 | 9.9 | 2.7 | 0 | 0.0 | 0 | 0 | 9 |
| A04 | 30.7 | 8 | 9.5 | 3.8 | 0 | 0.0 | 0 | 0 | 7 |
| A05 | 64.7 | 37 | 8.9 | 1.7 | 3 | 8.1 | 3 | 0 | 34 |
| A06 | 70.0 | 25 | 11.7 | 2.8 | 0 | 0.0 | 0 | 0 | 21 |
| A07_1 | 14.4 | 23 | 4.2 | 0.6 | 0 | 0.0 | 0 | 0 | 20 |
| A07_2 | 11.6 | 11 | 4.9 | 1.1 | 0 | 0.0 | 0 | 0 | 9 |
| A07_3 | 3.3 | 7 | 2.7 | 0.5 | 0 | 0.0 | 0 | 0 | 3 |
| A08_1 | 31.8 | 21 | 11.5 | 1.5 | 0 | 0.0 | 0 | 0 | 18 |
| A08_2 | 33.0 | 11 | 16.8 | 3.0 | 0 | 0.0 | 0 | 0 | 9 |
| A09 | 49.9 | 19 | 13.9 | 2.6 | 3 | 15.8 | 0 | 3 | 16 |
| A10_1 | 102.8 | 56 | 22.9 | 1.8 | 41 | 73.2 | 1 | 40 | 39 |
| A10_2 | 7.3 | 7 | 4.4 | 1.0 | 0 | 0.0 | 0 | 0 | 7 |
| B01_1 | 9.6 | 23 | 2.9 | 0.4 | 0 | 0.0 | 0 | 0 | 19 |
| B01_2 | 35.9 | 16 | 13.7 | 2.2 | 0 | 0.0 | 0 | 0 | 16 |
| B01_3 | 16.8 | 4 | 16.8 | 4.2 | 0 | 0.0 | 0 | 0 | 2 |
| B02 | 52.6 | 33 | 18.6 | 1.6 | 0 | 0.0 | 0 | 0 | 23 |
| B03 | 137.6 | 41 | 22.3 | 3.4 | 2 | 4.9 | 2 | 0 | 40 |
| B04_1 | 106.6 | 71 | 17.2 | 1.5 | 2 | 2.8 | 2 | 0 | 52 |
| B04_2 | 39.9 | 23 | 13.5 | 1.7 | 0 | 0.0 | 0 | 0 | 20 |
| B05 | 111.9 | 81 | 15.9 | 1.4 | 0 | 0.0 | 0 | 0 | 68 |
| B06 | 60.3 | 33 | 8.6 | 1.8 | 0 | 0.0 | 0 | 0 | 33 |
| B07 | 32.2 | 4 | 20.1 | 8.0 | 0 | 0.0 | 0 | 0 | 4 |
| B08 | 77.2 | 86 | 15.5 | 0.9 | 0 | 0.0 | 0 | 0 | 53 |
| B09 | 40.4 | 11 | 10.8 | 3.7 | 0 | 0.0 | 0 | 0 | 10 |
| B10 | 15.5 | 47 | 2.3 | 0.3 | 1 | 2.1 | 1 | 0 | 34 |
| Whole genome | 1414.8 | 855 | 12.4 | 2.2 | 52 | 3.8 | 9 | 43 | 689 |
| SDL: the number of segregation distorted loci on each linkage group (P<0.001). | | | | | | | | |  |
| SDL%: the percentage of distorted loci on each linkage group. | | | | | | |  |  |  |
| female parent: the number of loci distorted towards female parent Florida-07. | | | | | | | | |  |
| male parent: the number of loci distorted towards male parent GP-NC WS 16. | | | | | | | | |  |
| Whole genome: the total of all linkage groups was calculated for map distance, the number of loci, SDL, | | | | | | | | | |
| female and male parent. Average of linkage groups was calculated for largest gap, map density and SDL% | | | | | | | | | |
